# Direct inhibitors of InhA with efficacy similar or superior to isoniazid in novel drug regimens for tuberculosis

**DOI:** 10.1101/2024.03.08.584126

**Authors:** Lourdes Encinas, Si-Yang Li, Joaquin Rullas-Trincado, Rokeya Tasneen, Sandeep Tyagi, Heena Soni, Adolfo Garcia-Perez, Jin Lee, del Rio Rubén González, Jaime De Mercado, Verónica Sousa, Izidor Sosič, Stanislav Gobec, Alfonso Mendoza-Losana, Paul J. Converse, Khisi Mdluli, Nader Fotouhi, David Barros-Aguirre, Eric L. Nuermberger

**Author notes:** Address correspondence to Eric L. Nuermberger,; Lourdes Encinas. Alfonso Mendoza-Losana, Department of Bioengineering and Aerospace Engineering, Carlos III University of Madrid, 28040 Madrid, Spain.

## Abstract

Isoniazid is an important first-line medicine to treat tuberculosis (TB). Isoniazid resistance increases the risk of poor treatment outcomes and development of multidrug resistance, and is driven primarily by mutations involving *katG*, encoding the pro-drug activating enzyme, rather than its validated target, InhA. The chemical tractability of InhA has fostered efforts to discover direct inhibitors of InhA (DIIs). During the past five years, successful target engagement and *in vivo* efficacy have been demonstrated by diverse DIIs. In this study, we bridge the gap in understanding the potential contribution of DIIs to novel combination regimens and demonstrate a clear distinction of DIIs, like GSK693 and the newly described GSK138, from isoniazid, based on activity against clinical isolates and contribution to novel drug regimens. The results presented increase the understanding of DII mechanism of action and provide further impetus to continue exploiting InhA as a promising target for TB drug development.

## INTRODUCTION

Tuberculosis (TB) is a communicable disease caused by *Mycobacterium tuberculosis* (*M.tb*). Globally, an estimated 10.6 million people developed TB in 2022, up from best estimates of 10.3 million in 2021 and 10.0 million in 2020. Until the coronavirus (COVID-19) pandemic, TB was the leading cause of death from a single infectious agent, ranking above HIV/AIDS. The COVID-19 pandemic has had a negative impact on access to TB diagnosis and treatment as well as the burden of TB disease. The estimated number of deaths from TB increased between 2019 and 2021, reversing years of decline. An estimated total of 1.3 million people died from TB in 2022 (including 167,000 people with HIV). The net reduction in the global number of deaths caused by TB from 2015 to 2022 was 19%, far from the WHO End TB Strategy milestone of a 75% reduction by 2025 (1, 2). With timely diagnosis and treatment with first-line drugs, most people who develop TB are cured and onward transmission of infection is curtailed. The currently recommended treatment for drug-susceptible pulmonary TB is a 6-month regimen consisting of an *intensive phase* of 2 months with a 4-drug regimen of isoniazid, rifampicin, pyrazinamide, and ethambutol, followed by a *continuation phase* of four months with isoniazid and rifampicin.

Isoniazid was discovered in 1952 and has been widely used to treat TB ever since (3). It is a prodrug that requires activation by the mycobacterial catalase-peroxidase enzyme KatG to form the reactive isonicotinyl acyl radical (4), which then forms a covalent adduct with the cofactor nicotinamide adenine dinucleotide (isoniazid-NAD adduct) (5). This adduct is the inhibitor of the mycobacterial fatty acid synthase II (FAS-II) component enoyl-acyl carrier protein reductase (InhA), which is required for the synthesis of mycolic acids, a central component of the mycobacterial cell wall (6, 7).

Isoniazid is an important first-line TB drug. Baseline isoniazid resistance increases the risk of poor treatment outcomes (e.g. treatment failure or relapse) and acquisition of multidrug-resistant (MDR) TB. Based on evidence reviews indicating reduced efficacy of the standard first-line drugs for the treatment of isoniazid-resistant TB (Hr-TB) (8-13), the World Health Organization (WHO) issued a Supplement to its guidelines for the treatment of drug-resistant TB in 2018, providing new recommendations for the management of Hr-TB (14). Resistance to isoniazid is primarily caused by mutations in the activating enzyme KatG or in the upstream promoter region of InhA or more rarely in the InhA enzyme itself. Combinations of these mutations may also occur. By and large, the most common mutations in Hr-TB strains are found in *katG* and confer “high-level” resistance, even in the absence of an *inhA* mutation. In this situation, the inclusion of isoniazid in the regimen, even at high doses, is unlikely to increase its effectiveness (although this question is currently under investigation: *ClinicalTrials.gov Identifier: NCT01936831*). On the other hand, mutations in the *inhA* promoter or in the *inhA* gene are generally associated with lower-level resistance than *katG* mutations, and higher doses of isoniazid (10-15 mg/kg/day) may result in bactericidal activity against such *inhA* mutants similar to that observed with standard isoniazid doses (4-6 mg/kg/day) against fully susceptible strains (4, 15-17).

The opportunity to overcome the high rate of clinical resistance to isoniazid due to *katG* mutations, together with the biological relevance of InhA (target validated clinically by isoniazid and ethionamide) and its chemical tractability, (18) has fostered efforts to discover direct inhibitors of InhA (DIIs). During the last five years, three structurally different molecules have demonstrated *in vivo* efficacy in murine TB models upon oral administration: **NITD-916** (19), **GSK2505693A (GSK693)** (20) and **AN12855** (21).

## RESULTS

To our knowledge, **GSK693** was the first DII compound to demonstrate *in vivo* efficacy comparable to that of isoniazid (20). More recently, Xia and coauthors reported the discovery of a direct, cofactor-independent inhibitor of InhA, AN12855, which showed good efficacy in acute and chronic murine TB models that was also comparable to isoniazid (20). The high preliminary human dose prediction of **GSK693** hampered its further development as a lead compound. Within the same thiadiazole-based series, **GSK3081138A (GSK138),** a structurally very similar and slightly more lipophilic compound, was selected as a back-up compound based on its balanced profile of physicochemical properties, *in vitro* potency, *in vivo* pharmacokinetics (PK), and safety (Table 1).

**TABLE 1.**
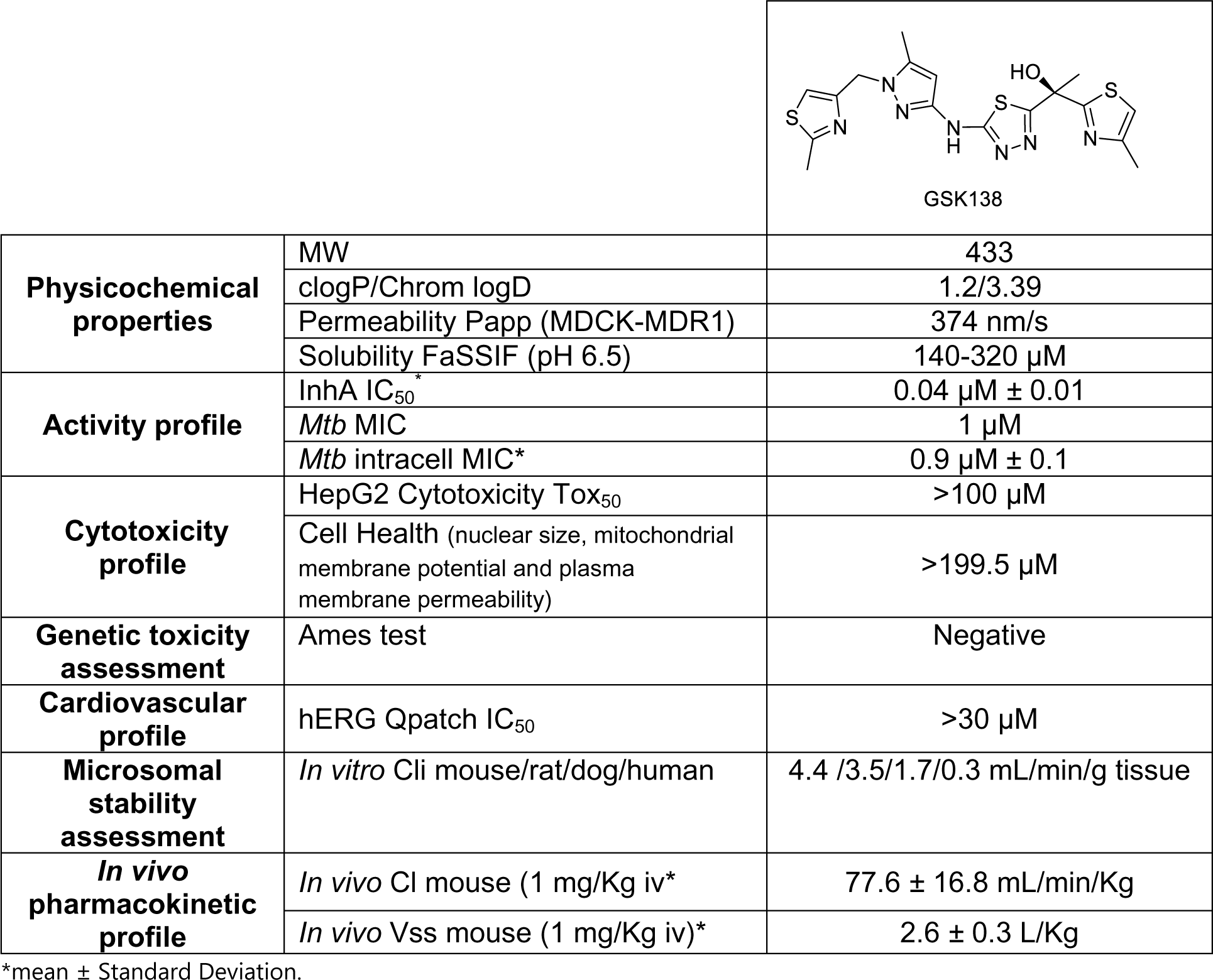
Structure and properties of the optimized lead **GSK138**. The lead was assessed for activity against *M. tuberculosis* H37Rv both intracellularly and extracellularly. The physicochemical and ADMET properties were determined as well.

|  |  |  |
| --- | --- | --- |
|  |  | GSK138 |
| **Physicochemical properties** | MW | 433 |
|  | clogP/Chrom logD | 1.2/3.39 |
|  | Permeability Papp (MDCK-MDR1) | 374 nm/s |
|  | Solubility FaSSIF (pH 6.5) | 140-320 µM |
| **Activity profile** | InhA IC<sub>50</sub>* | 0.04 µM ± 0.01 |
|  | *Mtb* MIC | 1 µM |
|  | *Mtb* intracell MIC* | 0.9 µM ± 0.1 |
| **Cytotoxicity profile** | HepG2 Cytotoxicity Tox<sub>50</sub> | >100 µM |
|  | Cell Health (nuclear size, mitochondrial membrane potential and plasma membrane permeability) | >199.5 µM |
| **Genetic toxicity assessment** | Ames test | Negative |
| **Cardiovascular profile** | hERG Qpatch IC<sub>50</sub> | >30 µM |
| **Microsomal stability assessment** | *In vitro* Cli mouse/rat/dog/human | 4.4 /3.5/1.7/0.3 mL/min/g tissue |
| ***In vivo* pharmacokinetic profile** | *In vivo* Cl mouse (1 mg/Kg iv)* | 77.6 ± 16.8 mL/min/Kg |
|  | *In vivo* Vss mouse (1 mg/Kg iv)* | 2.6 ± 0.3 L/Kg |

\*mean ± Standard Deviation.

**GSK138** is a medium molecular weight compound with a chrom logD at pH 7.4 of 3.38. The measured solubility in Fasted State Simulated Intestinal Fluid (FaSSIF) was high (Table 1). The permeability of **GSK138** predicted from Madin-Darby Canine Kidney (MDCK) cells was also high (Table 1). The efflux ratio determined by assays with and without incubation with a potent P-glycoprotein (P-gp) inhibitor indicated that it is a P-gp substrate.

**GSK138** inhibited recombinant InhA with an IC_50_ of 0.04 µM. The MIC was 1 µM against *M. tuberculosis* H37Rv and **GSK138** retained its activity against intracellular bacteria growing inside THP-1-derived macrophages *in vitro* (MIC 0.9 µM). Additionally, it showed no effect up to the highest concentration tested (200 µM) in the Cell Health assay (measuring membrane, nuclear, and mitochondrial damage). The preliminary toxicological profile showed an overall clean *in vitro* safety profile.

To assess the susceptibility of **GSK138** to P450-mediated phase I metabolism, metabolic stability was determined during incubation in CD1 mouse, Sprague Dawley rat, beagle dog, and human liver microsomes. **GSK138** exhibited moderate *in vitro* clearance in liver microsomes from the pre-clinical species, and low *in vitro* clearance in humans.

To determine the pharmacokinetic parameters, **GSK138** was administered intravenously (formulation: 5%DMSO/20%Encapsin in saline solution) as a single bolus dose in C57BL/6 mice at a target dose of 1 mg/Kg. All pharmacokinetic parameters were determined in whole blood (Table 1). A moderate clearance and a moderate volume of distribution were observed.

The minimum concentrations of **GSK693** and **GSK138** that inhibit 90% of isolates tested (MIC_90_) were determined against a set of drug-susceptible, multidrug-resistant (MDR) and extensively drug-resistant (XDR) *M.tb* clinical isolates. Both **GSK693** and **GSK138** retained activity against these clinical isolates (**GSK693** MIC_90_ = 1.87 μM; **GSK138** MIC_90_ = 3.75 μM), similar to the MICs against strain H37Rv in the same assay (Table 2). As expected for DIIs, the thiadiazole compounds have KatG-independent activity. No change in MIC was observed against isonazid-resistant clinical isolates carrying mutations in *katG* S315T. Clinical isolates carrying an *inhA* C-15T mutation have increased InhA production which confers low-level resistance to isoniazid. Among eight clinical isolates with the *inhA* C-15T mutation, three showed low-level resistance to the thiadiazoles (i.e., MICs for both DIIs ≥4 times the MIC against strain H37Rv). Thiadiazoles remained equally effective among the rest of the sensitive, MDR and XDR *M.tb* clinical isolates tested.

**TABLE 2.** Activity of **GSK693** and **GSK138** against resistant *M.tb* clinical isolates obtained from Vall d’Hebron Hospital, Barcelona. Resistance pattern: **H = isoniazid (*katG* S315T mutation:G, *inhA* promoter C-15T mutation:P)**, R = rifampicin, Z = pyrazinamide, M = moxifloxacin, T = ethionamide, S = streptomycin, E = ethambutol, K = kanamycin, A = amikacin, Cm = capreomycin, O = ofloxacin, Cp = ciprofloxacin, and Pas = para-aminosalicylic acid

| Strain # | Strain ID | Resistance | 693 MIC (µM) | 138 MIC (µM) |
| --- | --- | --- | --- | --- |
| 1 | 13243 | MOCp | <0.23 | <0.23 |
| 2 | 12569 | OCp | <0.23 | <0.23 |
| 3 | 9685A | H(G)RESZCpT | <0.23 | <0.23 |
| 4 | 7788 | RZCmK | <0.23 | 0.23 |
| 5 | 14388 | H(G)RESZCmAT | <0.23 | 0.47 |
| 6 | 13026 | H(G)RESCmKCpO | <0.23 | 0.47 |
| 7 | 13214 | H(P)SPas | <0.23 | 0.47 |
| 8 | 12733 | H(G)EZOTPas | <0.23 | 0.47 |
| 9 | 7957 | KCp | <0.23 | 0.47 |
| 10 | 10841 | **H(G)**RESZMO | <0.23 | 0.47 |
| 11 | 10071 | **H(G)**RESZKCp | <0.23 | 0.47 |
| 12 | 11881 | **H(G)**RESZCmKCp | <0.23 | 0.47 |
| 13 | 8059 | Cp | <0.23 | 0.47 |
| 14 | 14294 | ZMO | 0.23 | 0.23 |
| 15 | 11341 | **H(G)**REOPas | 0.23 | 0.47 |
| 16 | 13830 | **H(G)**RESZCmKMO | 0.47 | 0.47 |
| 17 | 14379 | **H(G)**RESZMO | 0.47 | 0.47 |
| 18 | 14883 | **H(G)**RESZCp | 0.47 | 0.94 |
| 19 | 13222 | SCmMCpO | 0.47 | 0.94 |
| 20 | 10492 | RSCp | 0.47 | 0.94 |
| 21 | 11586 | **H(G)**RESKCp | 0.94 | 1.875 |
| 22 | 11347 | **H(P)**RECmKO | 0.94 | 1.875 |
| 23 | 10027 | **H(G)**RESZCmKCp | 0.94 | 1.875 |
| 24 | 7786 | **H(P)**RESCmKCp | 1.875 | 1.875 |
| 25 | 10190 | **H(P)**REZCp | 1.875 | 1.875 |
| 26 | 7543 | **H(P)**RESZCmKCp | 1.875 | 1.875 |
| 27 | 11366 | **H(GP)**RESCmKO | 1.875 | 3.75 |
| 28 | 11348 | **H(P)**RESOTPas | 3.75 | 7.5 |
| 29 | 13229 | **H(P)**RZMCpOT | 7.5 | >15 |
| 30 | H37Rv | Susceptible (control) | <0.23 | 0.94 |
| 31 | 14639 | Susceptible (control) | 0.47 | 0.47 |

Based on **GSK138**’s overall profile, the therapeutic efficacy of **GSK138** against *M.tb* in an acute murine model of intratracheal infection was determined (see FIG 1).

**FIG 1.**
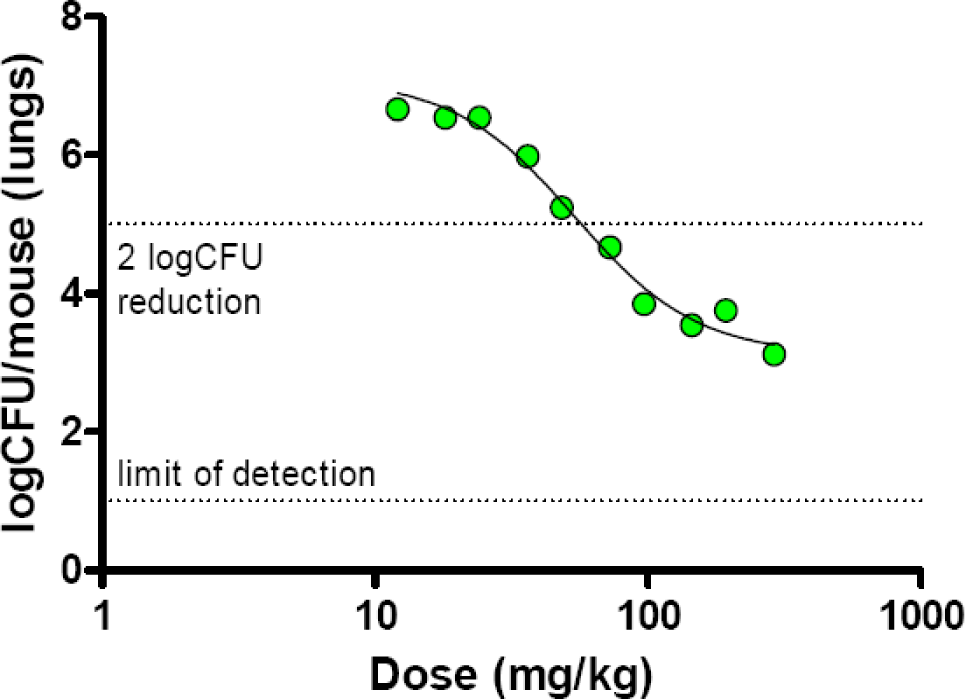
Dose-response relationship for **GSK138** in an acute mouse infection model of TB. Each point represents data from an individual mouse that received **GSK138** administered orally once daily for 8 days.

This acute infection model measures antitubercular activity on fast-growing bacteria (22). Treatment is administered for 8 consecutive days, starting 1 day after infection. Because the bacterial load reduction previously observed with **GSK693** was similar in both acute and chronic murine TB models (20), we performed a full dose-response study only in the acute model to characterize the compound and estimate the optimal dose for future combination studies. **GSK138** induced a net killing of the bacteria at the highest doses. The ED_99_ (the dose producing a 2-log_10_ reduction in colony-forming units [CFUs] compared to untreated control mice) for **GSK138** was 57 mg/Kg (95% confidence interval [CI]: 50-67 mg/Kg) and the dose of **GSK138** at which the 90% of the maximum bactericidal effect was achieved (ED_max_) was 167 mg/Kg (95%CI: 125->290 mg/Kg). The whole blood area under the concentration-time curve over 24 hours post-dose (AUC_0-24h_) at steady-state associated with this ED_max_ (AUC_EDmax_) was 68,544 ng*h/mL. Comparison with previous data suggests that **GSK138** is as efficacious as **GSK693** at a lower exposure, and therefore **GSK138** has the potential for a lower dose prediction in humans.

Ultimately, any antitubercular drug must be used in combination with other anti-tubercular drugs to treat active TB. The success of any new regimen will depend on the properties of these drugs and how they work in combination. **GSK693** and **GSK138** showed suitable profiles to justify investigation of the efficacy of these DIIs in combination with other drugs in animal models. Firstly, **GSK693** was selected as a tool compound to learn about the chemical series and its interactions with potential companion drugs. Experiment 1 was performed in a well-established high-dose aerosol infection model (23) with the following objectives: 1) to evaluate its ability to replace isoniazid (H) in combination with rifampicin (R) and pyrazinamide (Z) in the core first-line regimen, 2) to evaluate its ability to replace moxifloxacin (M) in combination with pretomanid (Pa) and pyrazinamide in the novel PaMZ regimen, and 3) to evaluate its contribution to the bactericidal activity of 2-drug combinations including bedaquiline (B), sutezolid (U), linezolid (L) and pretomanid (Figure 2).

**FIG 2.**
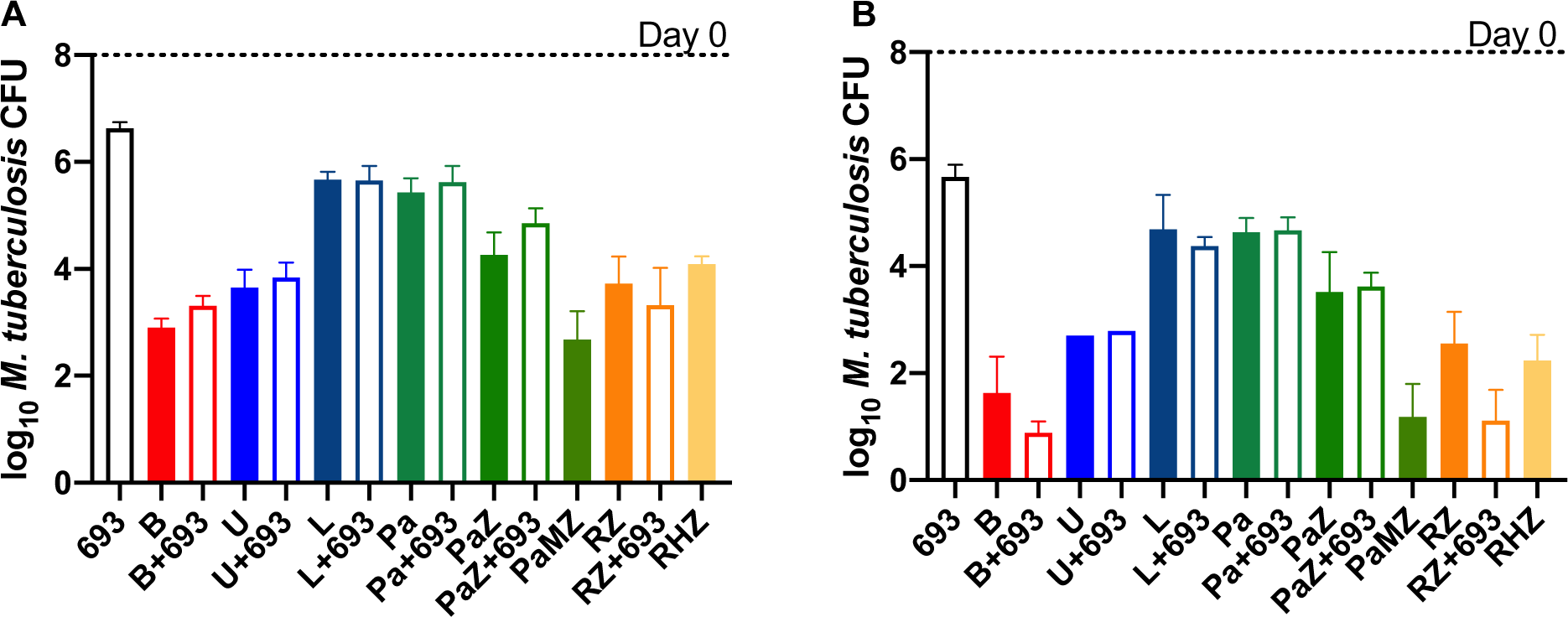
Mean (± SD) lung CFU counts at D0 and after 4 (A) or 8 (B) weeks of treatment in Expt 1. In combination with RZ, but not with other drugs, **GSK693** showed significantly enhanced antibacterial activity at week 8 (B) but not at week 4 (A). Open bars show lung CFU counts with the addition of **GSK693** to drugs shown in solid bars. Drug doses: R = rifampicin 10 mg/Kg, Z = pyrazinamide 150 mg/Kg, H = isoniazid 10 mg/Kg, 693 = **GSK693** 300 mg/Kg, Pa = pretomanid 50 mg/Kg, M = moxifloxacin 100 mg/Kg, B = bedaquiline 25 mg/Kg, L = linezolid 100 mg/Kg, U = sutezolid 50 mg/Kg.

In this model, in which untreated mice routinely succumb to infection with lung CFU counts above 8 log_10_ within the first 3-4 weeks after infection, **GSK693** (300 mg/Kg) reduced the lung CFU counts by 1.34 and 2.33 log_10_ after 4 and 8 weeks of treatment, respectively. This bactericidal effect approached that of linezolid or pretomanid. No additive effect was observed when **GSK693** was combined with sutezolid, linezolid or pretomanid, nor was it as effective as moxifloxacin in combination with pretomanid and pyrazinamide. However, the combination of **GSK693** with rifampicin and pyrazinamide (RZ) was significantly more active than RZ alone or in combination with isoniazid after 8 weeks of treatment (p<0.05). Notably, the combination of **GSK693** with bedaquiline also resulted in greater activity after 8 weeks of treatment (p=0.08) than that observed with bedaquiline alone. This additive effect of **GSK693** was attributable to its prevention of selection of bedaquiline-resistant mutants, as emergence of bedaquiline resistance was observed in 2 of the 4 mice treated with bedaquiline alone for 8 weeks, consistent with previous results (24). Excluding these 2 mice from the analysis revealed no difference between treatment with bedaquiline alone and bedaquiline plus **GSK693.**

The promising result observed with RZ+**GSK693** (in Exp. 1) prompted a follow-up experiment to confirm the additive effect of **GSK693**, evaluate the dose-response relationship for **GSK693** and explore potential drug-drug interactions in the RZ+**GSK693** combination.

As observed in Experiment 1, the addition of **GSK693**, but not isoniazid, significantly increased the activity of the RZ combination in Experiment 2 (Figure 3).

**FIG 3.**
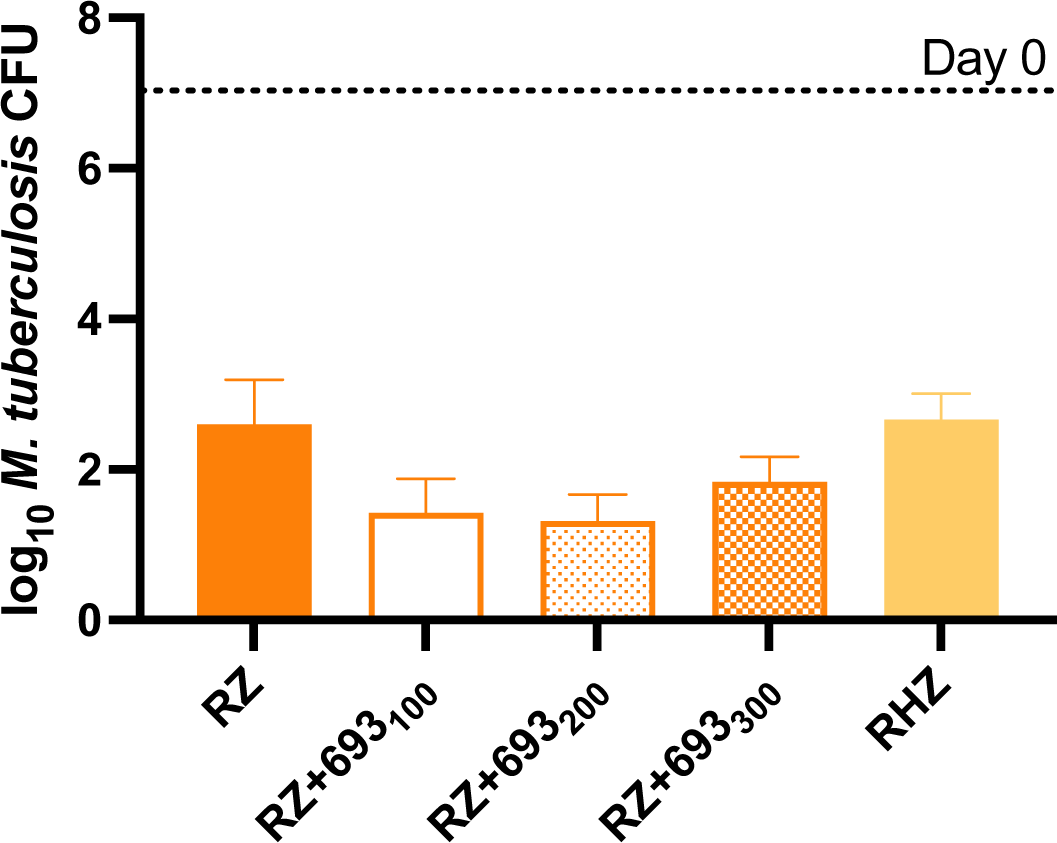
Mean (± SD) lung CFU counts at D0 and after 8 weeks of treatment in Experiment 2. 693 = **GSK693** significantly enhanced, in a non-dose-dependent manner, the activity of the RZ (rifampicin, 10 mg/Kg, plus pyrazinamide 150 mg/Kg) combination. Isoniazid (10 mg/Kg) did not enhance the activity of the combination. **GSK693** dose (in mg/Kg) is indicated in subscripts.

The magnitude of the additive effect was also similar between experiments. The addition of **GSK693** at 300 mg/Kg to RZ reduced the lung CFU counts by an additional 1.44 log in Experiment 1, as compared to a reduction of 0.76 log (p<0.05 vs RZ) in Experiment 2. Remarkably, however, greater reductions of 1.17 and 1.28 log (p<0.01 and 0.001 vs RZ, respectively) were observed when **GSK693** was used at 100 and 200 mg/Kg, respectively, in Experiment 2.

Although the sparse sampling prevented a formal assessment of the PK profile, the apparent lack of **GSK693** dose response was not explained by the **GSK693** exposures at the 100, 200 and 300 mg/Kg doses (Table 3).

**TABLE 3.**
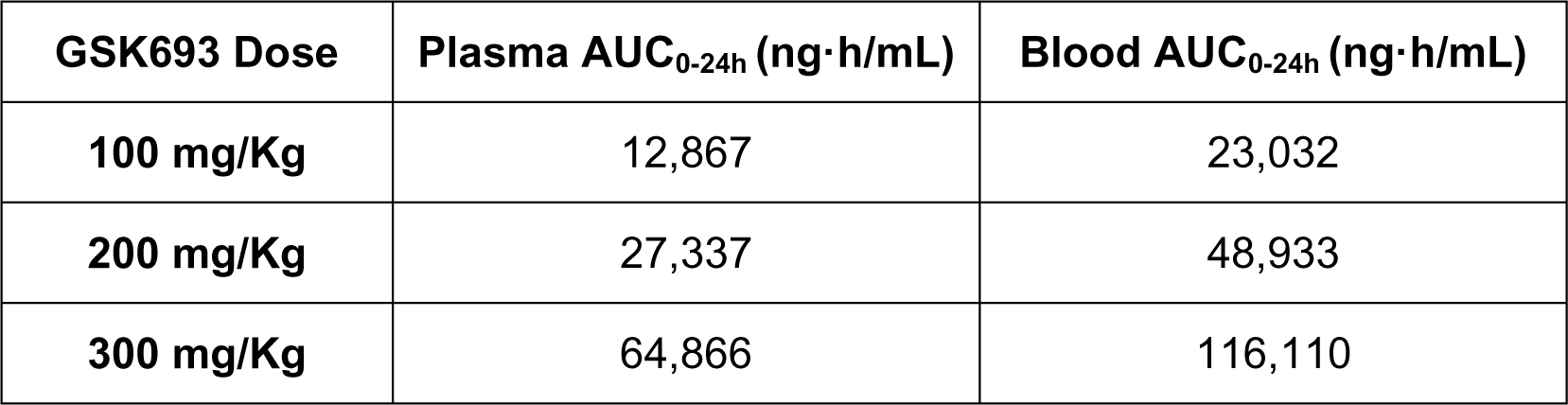
Data obtained from monocompartmental model of sparse plasma sampling concentrations in the combination study. A blood/plasma ratio of 1.79 was used to transform the plasma parameters to blood values.

| **GSK693 Dose** | **Plasma AUC<sub>0-24h</sub> (ng·h/mL)** | **Blood AUC<sub>0-24h</sub> (ng·h/mL)** |
| --- | --- | --- |
| **100 mg/Kg** | 12,867 | 23,032 |
| **200 mg/Kg** | 27,337 | 48,933 |
| **300 mg/Kg** | 64,866 | 116,110 |

Interestingly, the observed effect of **GSK693** in combination was achieved at a lower exposure than that needed to achieve the maximum effect in monotherapy in the acute infection model (110,200 ng·h/mL). Based upon the potential for drug-drug interactions, rifampicin was administered 1 hour prior to other drugs (25). Plasma exposure of rifampicin (AUC_0-24h_ = 68,544 ng·h/mL) when co-administered with pyrazinamide showed no evidence of a higher exposure that could explain the increase in the efficacy of the combination when compared to prior data for rifampicin when co-administered with pyrazinamide (AUC_0-24h_ = 160,600 ng·h/mL) (25) or as monotherapy at 10 mg/kg (AUC_0-24h_ = 87,200 to 142,100 ng·h/mL) (26).

The result from the combination of **GSK693** with RZ proved to be superior to the first-line treatment (RHZ). This result encouraged further combination experiments, now with **GSK138**.

A major objective of Experiment 3 was to determine the effect of adding **GSK138** to the novel regimen of bedaquiline, pretomanid, and linezolid (BPaL) recently approved for treatment of XDR and treatment-intolerant or non-responsive MDR TB and the effect of substituting **GSK138** for either bedaquiline or linezolid. The experiment also included the novel LeuRS inhibitor GSK3036656 (GSK656) (27, 28) that is now in phase 2 clinical trials. The objectives of this experiment were the following: 1) to evaluate the contribution of **GSK138** to the efficacy of 3- and 4-drug combinations based on the BPa backbone, and 2) to evaluate the effect of adding **GSK138** to the combination of rifampicin plus GSK656, with or without pyrazinamide (Figure 4).

**FIG 4.**
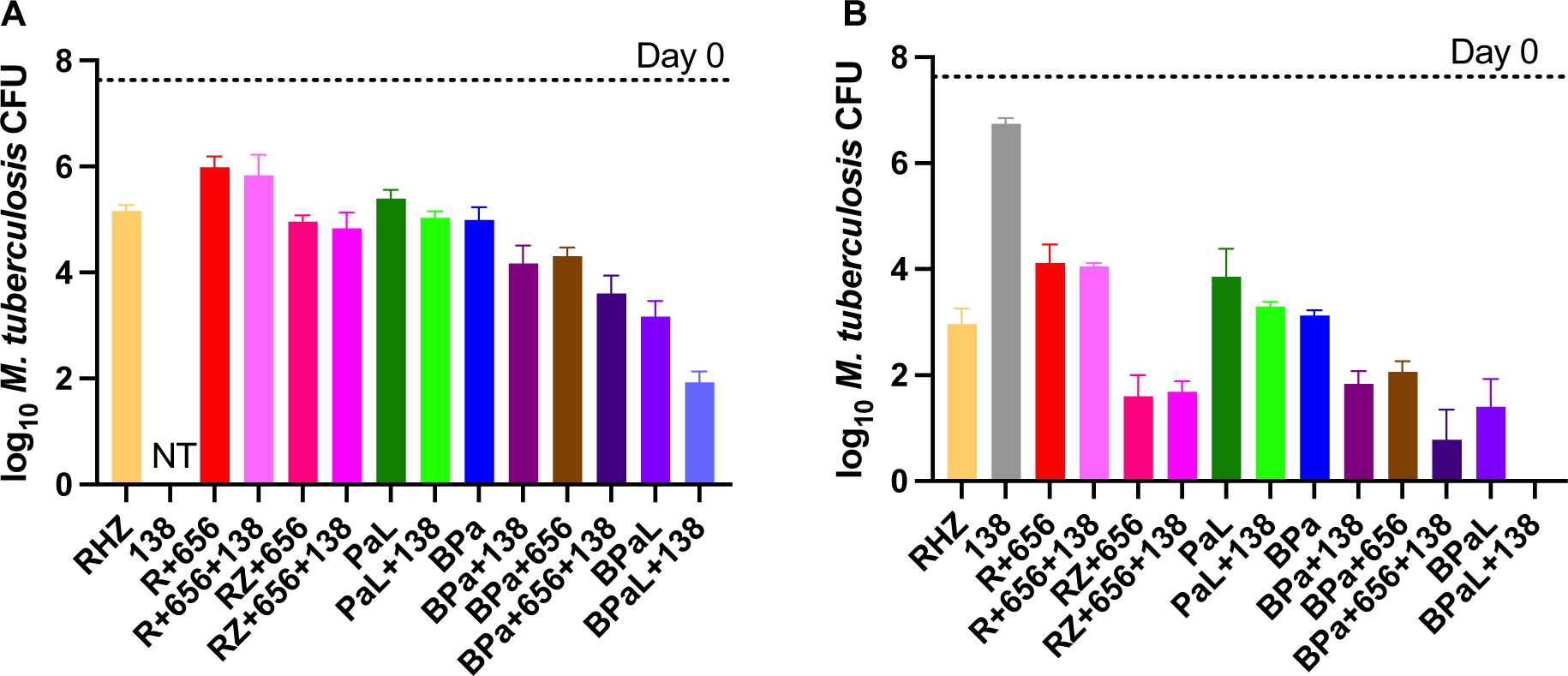
The direct InhA inhibitor **GSK138** enhanced the activity of the BPa, BPaL and BPa+GSK656 combinations after 4 weeks (A) or 8 weeks (B) of treatment. After 8 weeks of treatment, the BPaL+**GSK138** regimen rendered mouse lungs culture negative. Data are presented as mean (± SD) lung CFU counts. R = rifampicin 10 mg/Kg, Z = pyrazinamide 150 mg/Kg, H = isoniazid 10 mg/Kg, 138 = **GSK138** 200 mg/Kg, 656 = GSK656 (sulfate salt) 10 mg/Kg, Pa = pretomanid 50 mg/Kg, B = bedaquiline 25 mg/Kg, L = linezolid 100 mg/Kg. NT = not tested.

The addition of **GSK138** to BPaL, its BPa backbone, or the novel BPa+GSK656 regimen significantly increased the activity of each combination after 4 weeks (p<0.01) and after 8 weeks (p<0.0001) of treatment. Indeed, the 4-drug combination of BPaL plus **GSK138** was the only regimen tested to render all mice culture-negative after 8 weeks of treatment. After 8 weeks of treatment, the activity of the 3- and 4-drug regimens containing BPa plus **GSK138**, with or without GSK656, were statistically indistinguishable from that of BPaL and significantly superior to the first-line RHZ regimen (p<0.0001). The addition of **GSK138** did not significantly increase the activity of PaL, R+GSK656, or RZ+GSK656.

Experiment 4 (Figure 5) was performed to confirm the contribution of **GSK138** to the BPa backbone, this time including a range of **GSK138** doses. The experiment also included isoniazid as a comparator and combinations with GSK656 and the novel cholesterol-dependent inhibitor GSK2556286 (GSK286) (29) which is currently being investigated in a first time in human (FTIH) study to evaluate its safety, tolerability, and pharmacokinetics (NCT04472897).

**FIGURE 5.**
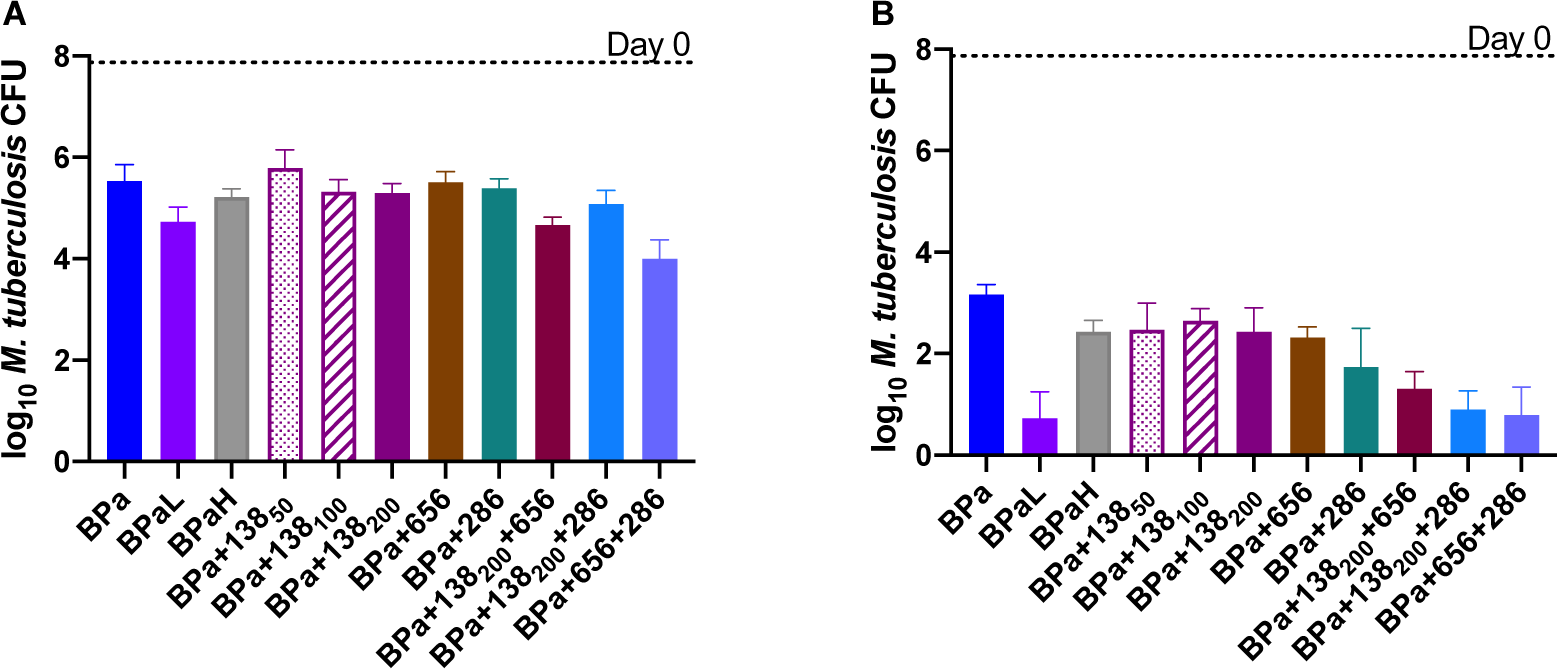
GSK138. significantly enhanced the activity of BPa and BPa-based regimens at 4 weeks (A) or 8 weeks (B), particularly in combination with GSK286. In combination with GSK656 or GSK286, the 200 mg/Kg dose of **GSK138** was used. Data are presented as mean (± SD) lung CFU counts B = bedaquiline, 25 mg/Kg; Pa = pretomanid, 100 mg/Kg, L = linezolid, 100 mg/Kg, H = isoniazid 10 mg/Kg, 286 = GSK286, 50 mg/Kg, 656 = GSK656 (hydrochloride salt), 10 mg/Kg. **GSK138** dose (in mg/Kg) is indicated in subscripts.

The results reaffirmed the additive effects of **GSK138** when added to BPa for 8 weeks of treatment, and similar results were observed with the addition of isoniazid or GSK656 to BPa. No dose-response of **GSK138** was evident after 8 weeks; and unlike in Experiment 3, BPa plus **GSK138** was less effective than BPaL (p<0.01 after 4 and 8 weeks). However, as observed in Experiment 3, the additive 4-drug combination of BPa+GSK656 with **GSK138** was statistically indistinguishable from BPaL, as was the combination of BPa+GSK286 with **GSK138**. These 4-drug combinations of BPa+**GSK138** plus either GSK656 or GSK286 also had bactericidal activity similar to BPa+GSK656+GSK286 after 8 weeks of treatment.

Given the superior bactericidal activity of the BPaL plus **GSK138** regimen compared to BPaL alone in Experiment 3, Experiment 5 was performed to determine whether addition of GSK138 to BPaL could shorten the duration of treatment needed to prevent relapse (Figure 6). Comparator regimens included BPaL plus one of the following: isoniazid, NITD-113 (prodrug for NITD-916, a previously reported DII based on a different scaffold than **GSK138**) (19) and moxifloxacin (M). As observed in Experiment 3, the addition of **GSK138** at a dose of 200 mg/Kg significantly increased the bactericidal activity of BPaL after 4 weeks of treatment (p<0.01), as did isoniazid (p<0.01), while there was a trend towards enhanced activity with NITD-113 (p=0.11) and moxifloxacin (p=0.10). BPaL+**GSK138** resulted in fewer culture-positive mice and a lower mean CFU count after 8 weeks compared to BPaL and BPaL plus other InhA inhibitors, although these differences were not statistically significant. Only the BPaLM regimen rendered all mice culture-negative at this time point. Similarly, the addition of **GSK138**, isoniazid or moxifloxacin to BPaL each reduced the proportion of mice relapsing after 8 and 12 weeks of treatment compared to BPaL alone, although the differences were statistically significant only after 8 weeks of treatment with moxifloxacin or isoniazid, as shown in Figure 6C.

**FIG 6.**
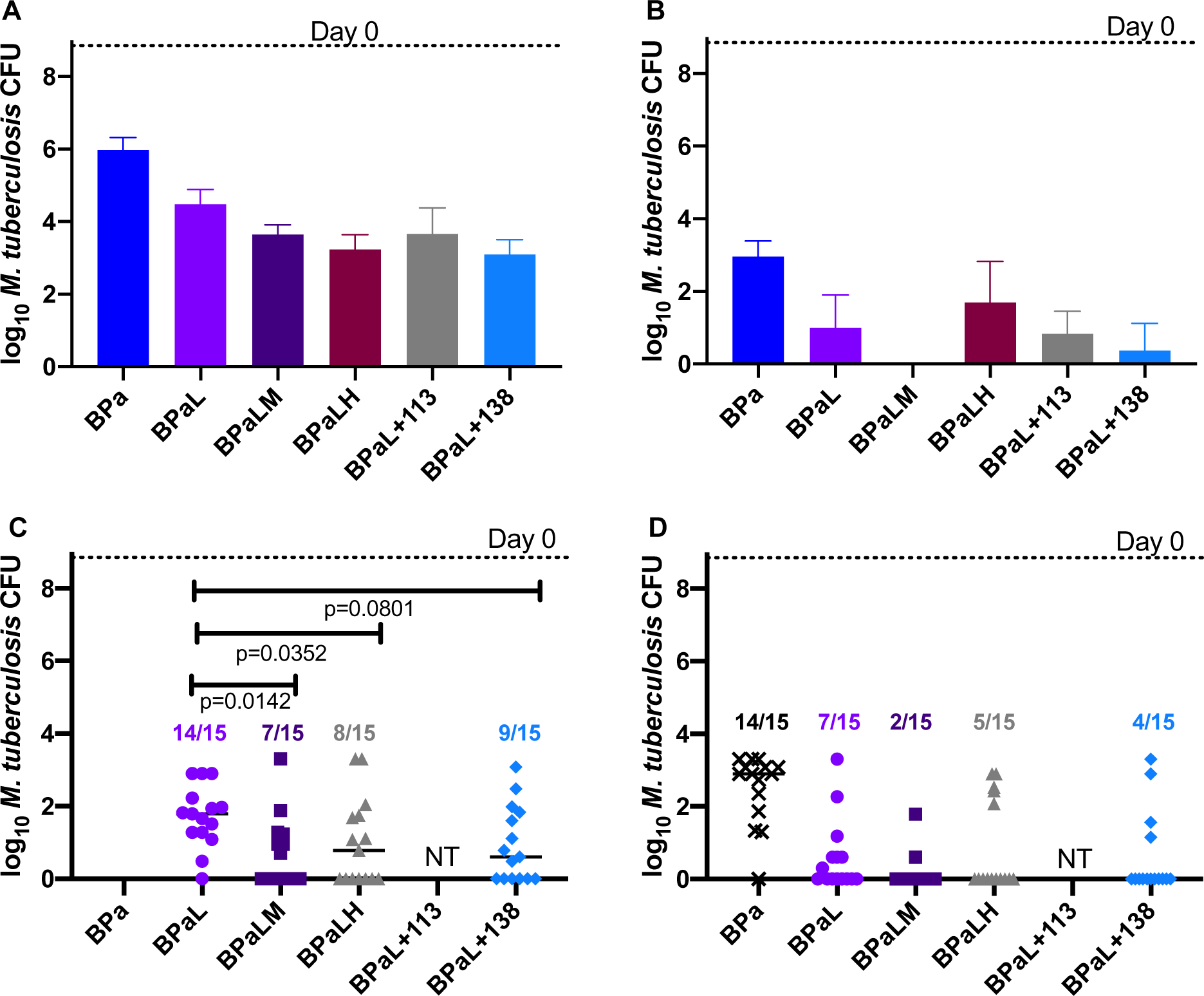
The addition of an InhA inhibitor or moxifloxacin enhanced the bactericidal and sterilizing activity of the BPaL regimen. After both 4 weeks (A) and 8 weeks (B), the addition of moxifloxacin, isoniazid, NITD-113, or **GSK138** to BPaL enhanced the bactericidal activity compared to BPaL alone (with the exception of isoniazid at week 8). Data are presented as mean (± SD) lung CFU counts. The proportion of mice relapsing after 8 weeks of treatment, followed by 12 weeks of no treatment (C) was statistically significantly lower in the presence of either moxifloxacin or isoniazid and approached statistical significance with **GSK138**. There were fewer relapses after 12 weeks of treatment and 12 weeks of follow-up (D) with the addition of a fourth drug, although these differences were not statistically significant. The proportions of mice relapsing are indicated above the symbols for lung CFU counts. B = bedaquiline, 25 mg/Kg, Pa = pretomanid, 100 mg/Kg, L = linezolid, 100 mg/Kg, M = moxifloxacin 100 mg/Kg, H = isoniazid 10 mg/Kg, NITD-113 = prodrug for NITD-916 (see Introduction) 150 mg/Kg, 138 = **GSK138**200 mg/Kg. NT = not tested.

## DISCUSSION

The thiadiazole-based DIIs (namely **GSK693**) proved capable of replacing isoniazid in the first-line regimen. In fact, the bactericidal activity of the regimen increased with this substitution and **GSK693** increased the activity of the rifampicin-pyrazinamide combination. These results suggest that use of a DII instead of isoniazid could also increase the sterilizing activity of the regimen. The superior activity of the DII in this regimen may be the result of superior killing of phenotypically INH-tolerant persisters that have relatively lower *katG* expression, whether stochastically or in response to stress (30). Further development of thiadiazole DIIs could yield superior first-line regimens containing rifamycins and pyrazinamide.

The thiadiazole-based DIIs (namely **GSK138**) also proved capable of increasing the bactericidal and sterilizing activity of BPa-based regimens. BPaL is an effective 6-month all-oral regimen for XDR-TB and difficult-to-treat MDR-TB cases (31, 32). The improved efficacy observed with the addition of **GSK138** suggests that this or another DII could further improve the BPaL regimen by increasing the overall cure rate, shortening treatment duration and/or reducing the emergence of drug resistance. Considering the strong overall activity of BPa+**GSK693** and **GSK693**’s ability to prevent selection of bedaquiline-resistant mutants, the DIIs of this class could also reduce the need for linezolid, the most toxic component of the BPaL regimen allowing lower doses and/or shorter durations of linezolid.

The use of thiadiazole DIIs alone or in combination with GSK656 to replace linezolid entirely, if proven safe, could enable the use of BPa-based regimens as alternative, more universally active, first-line regimens that would be less affected by isoniazid monoresistance or MDR.

Although not the focus of this report, we observed that the addition of moxifloxacin to BPaL improved the bactericidal and sterilizing activity of the regimen. BPaL and BPaLM were studied in the TB-PRACTECAL trial (ClinicalTrials.gov Identifier: NCT02589782). Our results, which were obtained before the start of the trial, predicted superior efficacy of the 4-drug combination. Indeed, a higher rate of sputum culture conversion at 8 weeks was observed in TB-PRACTECAL participants receiving treatment with BPaLM vs. BPaL (77% vs. 46%) (33).

This research adds to the limited knowledge of the activity of direct InhA inhibitors in combination with new and existing TB drugs. The results suggest that a direct InhA inhibitor (e.g., **GSK138** and **GSK693**) could be a promising partner in novel drug regimens, enhancing their efficacy and/or preventing the selection of bedaquiline-resistant mutants. These findings increase our understanding of the mechanism of action of direct inhibitors of InhA and provide further impetus to continue exploiting InhA as a promising target for TB drug development.

## MATERIALS AND METHODS

The human biological samples were sourced ethically, and their research use was in accord with the terms of the informed consents under an IRB/EC approved protocol. All animal studies performed by GSK were ethically reviewed and carried out in accordance with European Directive 2010/63/EEC and the GSK Policy on the Care, Welfare and Treatment of Animals. All animal studies performed at Johns Hopkins University (JHU) were conducted in accordance with the GSK Policy on the Care, Welfare and Treatment of Laboratory Animals and were approved by the Institutional Animal Care and Use Committee of JHU.

### Chemistry

A micromilling method was applied to **GSK138** for particle size reduction in order to obtain a micronized GSK138 that was used in *in vivo* experiments. The Mixer Mill MM 301 (Retsch) was used at a frequency of 20 Hz for four cycles of 5 minutes.

NMR spectra were recorded on an Agilent Inova 600 MHz spectrometer equipped with a 5 mm Triple Resonance Gradient Probe IDTG600-5 (experiments run under software version vnmr3.2 Revision A). Measurements were made at room temperature in DMSO-d6 solvent. The chemical shift (d) values are expressed in parts per million (ppm) and coupling constants are in Hertz (Hz). The chemical shifts (δ) were given relative to the residual ^1^H and ^13^C signals of the solvent peak as an internal standard: in ^1^H NMR (600 MHz) δ 2.49 ppm (quin, C_2_D_5_HOS) for DMSO-*d_6_*; in ^13^C NMR (150 MHz) δ 40.07 ppm (sept) for DMSO-*d_6_*. Legend: s = singlet, d = doublet, sept = septet, br = broad signal. LC-MS purity data were collected using a Waters Acquity UPLC instrument coupled with Waters Acquity single quadrupole mass and photodiode array detectors. High-resolution MS (HRMS) was performed on a QSTAR Elite System mass spectrometer. ^1^H NMR (600 MHz, DMSO-*d*_6_): δ 10.67 (s, 1H, NH), 7.26 (s, 1H, OH), 7.18 (s, 1H), 7.09 (s, 1H), 5.79 (s, 1H), 5.16 (s, 2H), 2.59 (s, 3H), 2.29 (s, 3H), 2.26 (s, 3H), 1.97 (s, 3H). ^13^C NMR (150 MHz, DMSO-*d*_6_): δ 175.96, 166.46, 166.29, 164.69, 152.55, 152.26, 147.61, 140.88, 116.47, 115.38, 94.56, 74.68, 48.82, 28.98, 19.29, 17.51, 11.53. HRMS (ESI) *m*/*z*: calcd for C_17_H_19_N_7_OS_3_ [M + H]^+^, 434.0891; found, 434.0889.

### Permeability studies

Studies were performed as described by Polli et al. (34), with minor modifications. GF120918 was used as the inhibitor of P-gp. Apical-to-basolateral (A-to-B) and basolateral-to-apical (B-to-A) transport were studied across MDR1-MDCKII cell monolayers in the absence and presence of the P-gp inhibitor GF120918, and the Papp (intrinsic apparent permeability) was estimated in both directions with or without inhibitor.

### Solubility studies

Solubility assays were performed using a miniaturized shake flask method. 10 mM stock solutions of each test compound were used to prepare calibration standards (10-220 μM) in DMSO, and to spike (1:50) duplicate aqueous samples of FaSSIF (simulating fasting state biorelevant media, pH 6.5), with a final DMSO concentration of 2%. After shaking for 2 hours at 25 °C, the solutions were filtered and analysed by means of HPLC-DAD (Agilent 1200 Rapid Resolution HPLC with a diode array detector). Best fit calibration curves were constructed using the calibration standards, which were used to determine the aqueous samples solubility (35).

### Bacterial strains

*M. tuberculosis* H37Rv was mouse-passaged, frozen in aliquots and sub-cultured in Middlebrook 7H9 broth with 10% oleic acid-albumin-dextrose-catalase (OADC) (Fisher, Pittsburgh, PA) and 0.05% Tween 80 prior to high-dose mouse aerosol infection. MDR and XDR *M. tuberculosis* clinical isolates representing different resistance phenotypes belong to the collection of strains of the Vall d’Hebron hospital of Barcelona.

*M. tuberculosis* H37Rv and H37Rv-Luc were routinely propagated at 37°C in Middlebrook 7H9 broth (Difco) supplemented with 10% Middlebrook albumin-dextrose-catalase (ADC)(Difco), 0.2% glycerol and 0.05% (vol/vol) tyloxapol or on Middlebrook 7H10 agar plates (Difco) supplemented with 10% (vol/vol) OADC (Difco). Hygromycin B was added to the medium (50 μg/mL) to ensure plasmid maintenance when propagating the H37Rv-Luc strain. This strain constitutively expresses the luciferase *luc* gene from *Photinus pyralis* (GenBank Accession Number M15077) cloned in a mycobacterial shuttle plasmid derived from pACE-1 (36).

### Intracellular MIC assay

Frozen stocks of macrophage THP-1 cells (ATCC TIB-202) were thawed in RPMI-1640 medium (Sigma) supplemented with 10% fetal bovine serum (FBS) (Gibco), 2 mM L-glutamine (Sigma) and 1 mM sodium pyruvate (Sigma). THP-1 cells were passaged only five times and maintained without antibiotics between 2–10 × 10^5^ cells/mL at 37 °C in a humidified, 5% CO2 atmosphere. THP-1 cells (3 × 10^8^) were simultaneously differentiated with phorbol myristate acetate (PMA, 40 ng/mL, Sigma) and infected for 4 hours at a multiplicity of infection (MOI) of 1:1 with a single cell suspension of H37Rv-Luc. After incubation, infected cells were washed four times to remove extracellular bacilli and resuspended (2 × 10^5^ cells/mL) in RPMI medium supplemented with 10% FBS (Hyclone), 2 mM L-glutamine and pyruvate and dispensed in white, flat bottom 384-well plates (Greiner) in a final volume of 50 μL (max. 0.5% DMSO). Plates were incubated for 5 days under 5% CO_2_ atmosphere, 37 °C, 80% relative humidity before growth assessment. The Bright-Glo™ Luciferase Assay System (Promega, Madison, WI) was used as cell growth indicator for the H37Rv-Luc strain. Luminescence was measured in an Envision Multilabel Plate Reader (PerkinElmer) using the opaque 384-plate Ultra Sensitive luminescence mode, with a measurement time of 50 ms. A 90% reduction in light production was considered growth inhibition and the IC_90_ value was interpolated from the dose response curve.

### Extracellular MIC assays

MICs against the H37Rv strain were determined by broth dilution assay in Middlebrook 7H9 medium supplemented with 10% ADC. After incubating at 37 °C for six days, 25 µL resazurin solution (one tablet in 30 mL sterile PBS) was added to each well. Following incubation at 37 °C for two additional days, the lowest concentration of drug that inhibited 90% of resazurin conversion compared to internal DMSO control wells with no drug added was used to define MIC values.

MICs against clinical isolates of *M. tuberculosis* were determined using the mycobacteria growth indicator tubes (MGIT) system. Approximately 1 mg wet weight from a Lowenstein-Jensen slant, with an estimated bacterial load of 10^8^ CFU/mL, was inoculated into McCartney vials containing 1 mL of distilled water and 5 glass beads. The mixtures were homogenized by vortexing for 1-3 minutes. The opacity of the suspensions was adjusted by the addition of sterile distilled water to that of a 0.5 McFarland turbidity standard. 100 µL were used to inoculate MGIT vials containing serial dilutions of the compounds. MIC values were defined using the BACTEC MGIT 960 System (Becton Dickinson) and following the manufacturer’s instructions.

### HepG2 cytotoxicity assay

HepG2 cells were cultured using Eagle’s minimum essential media (EMEM) supplemented with 10% heat-inactivated FBS, 1% Non-Essential Amino Acid (NEAA), and 1% penicillin/streptomycin. Prior to addition of the cell suspension, 250 nL of test compounds per well were predispensed in Tissue culture -treated black clear-bottomed 384-well plates (Greiner, cat. no. 781091) with an Echo 555 instrument. After that, 25 μL of HepG2 (cat. no. ATCC HB-8065) cells (∼3000 cells/well) grown to confluency in EMEM supplemented with 10% heat-inactivated FBS, 1% NEAA, and 1% penicillin/streptomycin were added to each well with the reagent dispenser. Plates were allowed to incubate at 37 °C with 20% O_2_ and 5% CO_2_ for 48 hrs. After incubation, the plates were equilibrated to room temperature before ATP levels were measured with the CellTiter Glo kit (Promega) as the cell viability read-out. 25 μL of CellTiter Glo substrate dissolved in the buffer was added to each well. Plates were incubated at room temperature for 10 min for stabilization of luminescence signal and read on a View Lux luminometer with excitation and emission filters of 613 and 655 nm, respectively. The Tox_50_ value corresponds to the concentration of the compound necessary to inhibit 50% of cell growth.

### Cell Health Assay

This is a 3-parameter automated imaging cell-based assay to measure the cytotoxic effect of compounds in human liver-derived HepG2 cells. Using fluorescent staining, the key parameters measured in this assay are nuclear size, mitochondrial membrane potential and plasma membrane permeability. HepG2 cells (ATCC HB-8065) were incubated with the test compounds in 384-well plates. After 48 hours, the staining cocktail was added. Hoechst 33342 is used to stain nuclei and quantify changes in nuclear morphology. Tetramethylrhodamine, methyl ester (TMRM) is a cationic dye that accumulates in healthy mitochondria that maintain a mitochondrial membrane potential and leaks out of mitochondria when the mitochondrial membrane potential is dissipated. TOTO-3 iodide labels nuclei of permeabilized cells and is used to measure plasma membrane permeability. Following 45 min of incubation with these stains, the plates were sealed using a black seal for reading on an INCell Analyzer 2000 (GE Healthcare). Each parameter measurement produces the percentages of cells which are ‘LIVE’ or ‘DEAD’. The IC_50_ is defined as the compound concentration that inhibits 50% of cell growth.

### Ames Assay

The Ames assay was carried out as previously described (37) using all strains.

### hERG Assay

hERG activity was measured as previously described (38).

### Hepatic microsome stability

Human and animal microsomes and compounds were preincubated at 37 °C prior to addition of NADPH to final protein concentration of 0.5 mg/mL and final compound concentration of 0.5 µM. Quantitative analysis was performed using specific LC-MS/MS conditions. The half-life, elimination rate constant, and intrinsic clearance (mL/min/g tissue) were determined. The well-stirred model was used to translate to *in vivo* Cl values (mL/min/Kg).

### *In vivo* pharmacokinetics analysis

Single-dose pharmacokinetics experiments were performed in female C57BL/6 mice, 21-29 g, obtained from Charles River Laboratories (Wilmington, MA) and housed in cages in groups of three animals with water and food *ad libitum*. Animals were maintained for one week before the experiment.

The compound was dissolved in 20% Encapsin (Sigma-Aldrich), 5% DMSO (Sigma Aldrich) in saline solution (Sigma Aldrich) for intravenous administration and in 1% methylcellulose (Sigma-Aldrich) in water for oral administration.

For PK analysis, 25 µL of tail blood were collected by microsampling at 0.08 h, 0.25 h, 0.5 h, 1 h, 2 h, 4 h, 6 h, 8 h and 24 h for intravenous pharmacokinetics and 0.25 h, 0.5 h, 1 h, 2 h, 4 h, 6 h, 8 h and 24 h for oral pharmacokinetics.

### Assessment of acute efficacy in murine TB models

Specific pathogen-free, 8-10 week-old (18-20 grams) female C57BL/6 mice were purchased from Harlan Laboratories and were allowed to acclimate for one week. The experimental design for the acute assay has been previously described (22). In brief, mice were intratracheally infected with 100,000 CFU/mouse of *M. tuberculosis* H37Rv. Compounds were administered daily for 8 consecutive days starting 24 hours after infection. Lungs were harvested on day 9. All lung lobes were aseptically removed, homogenized and frozen. Homogenates were thawed and plated on 7H11 medium supplemented with 10% OADC plus 0.4% activated charcoal to reduce the effects of compound carryover. CFU were counted after 18 days of incubation at 37 °C. Log_10_ CFU vs. dose data were plotted. A sigmoidal dose-response curve was fitted and used to estimate ED_99_ and ED_max_. Data were analyzed using GraphPad software (Prism). The ED_99_ was defined as the dose in mg/Kg that reduced the number of CFUs in the lungs of treated mice by 99% compared to untreated infected mice. The EDmax is the dose in mg/g that resulted in 90% of predicted maximal effect.

### Modelling and simulations

The calculated exposures at ED_max_ for **GSK138** and **GSK693** were obtained using the IV mouse PKs profiles fitting to a bicompartmental model to obtain those parameters to simulate the oral whole blood exposures at ED_max_. Additionally, a monocompartmental model was used to fit the experimental oral pharmacokinetic studies in non-infected mouse together with measured plasma concentrations obtained from the sparse sampling in the Experiment 2 in infected mice. Parameters obtained from this fitting were used to simulate the profile after **GSK693** administration at 100, 200 and 300 mg/Kg in the combination study and to calculate the associated exposures (see Table 3).

### Blood and Plasma pharmacokinetic sampling and analysis

#### Sample collection from non-infected animals (PK studies)

Blood samples (25 µL) were taken from the lateral tail vein using a micropipette and were mixed, vortexed with 25 µL of saponin 0.1% and frozen at -80 °C until analysis.

#### Sample collection from infected animals

Mouse tail vein blood was collected at the indicated time points. Briefly, an incision was made in the lateral tail vein. 20-50 µL of blood was collected in BD Vacutainer® PST™ lithium heparin tubes from each mouse. The tubes were kept on ice before being centrifuged at 8000 rpm for 5 minutes. The supernatant plasma (15-30 µL) was transferred to labeled microcentrifuge tubes, frozen and stored at -80 °C and then shipped on dry ice to GSK for further analysis.

#### Sample pretreatment and LC-MS/MS analysis

10 µL of plasma or blood samples thawed at ambient temperature was mixed with 200 µL of ACN:MeOH (80:20). After this protein precipitation step, samples were filtered using a 0.45 µm filter plate (Multiscreen Solvinert 0.45um FTPE, Millipore) and then filtered using a 0.2 µm filter (AcroPrep Advance 96 Filter Plate 350 μL, 0.2 μm PTFE) to ensure sterilization prior to LC-MS analysis.

An Acquity Ultra-Performance liquid chromatography (UPLC) system (Waters Corp., Milford, MA, USA) coupled to a triple quadrupole mass spectrometer (API 4000™, AB Sciex, Foster City, CA, USA) was used for the analysis.

The chromatographic separation was conducted at 0.4 mL/min in an Acquity UPLC™ BEH C18 column (50×2.1 mm i.d., 1.7 mm; Waters Corp.) at 40°C with acetonitrile (ACN) (SigmaAldrich) and 0.1% formic acid as eluents.

Sciex Analyst software was used for the data analysis. The non-compartmental data analysis (NCA) was performed with Phoenix WinNonlin software in order to determine pharmacokinetic parameters and exposure.

### High-dose aerosol mouse infection model

Female specific pathogen-free BALB/c mice, aged 5-6 weeks, were purchased from Charles River (Wilmington, MA). Mice were infected by aerosol using the Inhalation Exposure System (Glas-col, Terre Haute, IN) using a log phase culture of *M. tuberculosis* H37Rv with an OD_600_ of 0.8-1 to implant approximately 3.5-4 log_10_ CFU in the lungs. Treatment started 2 weeks later (D0). Mice were sacrificed for lung CFU counts the day after infection and on D0 to determine the number of CFU implanted and the number present at the start of treatment, respectively.

### Antibiotic treatment

Mice were treated with the drugs and drug combinations indicated in Figures 2 through 6 at the following doses (in mg/Kg body weight): bedaquiline (25), pretomanid (50 or 100), moxifloxacin (100), linezolid (100), isoniazid (10), rifampicin (10), sutezolid (50), **GSK138** (50, 100, or 200), **GSK693** (100, 200, or 300), GSK656 (10), GSK286 (50), NITD-113 (150), and pyrazinamide (150). **GSK693**, **GSK138**, and GSK286 were formulated in 1% methylcellulose solution. GSK656 was formulated in distilled water. Other drugs were formulated as previously described (39-41). Bedaquiline and pretomanid were administered in back-to-back gavages and separated from companion drugs by at least 3 hours. Rifampicin was administered alone at least one hour before any companion drug.

### Evaluation of drug efficacy

Efficacy determinations were based on lung CFU counts after 4 or 8 weeks of treatment and, in one experiment, cohorts of mice were also kept for 12 weeks after completing 8 or 12 weeks of treatment to assess for relapse-free cure. At each time point, lungs were removed aseptically and homogenized in 2.5 mL of PBS. Serial 10-fold dilutions of lung homogenate were plated on selective 7H11 agar plates. To assess for relapse-free cure, the entire lung homogenate was plated. In experiments with bedaquiline, lung homogenates were plated on 7H11 agar supplemented with 0.4% activated charcoal to reduce drug carryover and doubling the concentrations of selective antibiotics in the media to mitigate binding to charcoal.

### Statistical analysis

Group means were compared by one-way ANOVA with Dunnett’s correction for multiple comparisons or by Student’s t-test, as appropriate, using GraphPad Prism version 8.

## ACKNOWLEDGEMENTS

We acknowledge funding by the European Union Seventh Framework Programme (FP7/2007-2013) under grant agreement N° 261378. We acknowledge Kala Barnes-Boyle for her technical assistance, Eva María López-Román and María José Rebollo-López for the MIC determination against mycobacteria and clinical isolates respectively, Raquel Gabarró, Jesús Gómez, Douglas J. Minick for conducting structural characterization experiments, Pablo Castañeda-Casado for the safety assessment and Fatima Ortega-Muro for her contribution in the review of the ADME and pharmacokinetics.

